# Estradiol induces global changes in miRNA expression in endometrial cancer cells and upregulates oncogenic miR-182

**DOI:** 10.1101/2024.05.12.593753

**Authors:** Klaudia Klicka, Tomasz M. Grzywa, Jarosław Wejman, Joanna Ostrowska, Paweł K. Włodarski

## Abstract

Endometrial cancer (EC) occurs mainly in perimenopausal age. Risk factors are associated with unopposed estrogen stimulation of the endometrium. There are two main types of EC, EC1 and EC2. The pathogenesis of EC1 is estrogen-dependent. MiRNAs are small RNAs that regulate gene expression posttranscriptionally. They are induced by estrogens in different hormone-dependent neoplasias including breast cancer. However, little is known regarding their role in EC. Thus, this study aims to assess the role of the miRNA-estrogen axis in endometrial cancer cells. Estrogen-dependent endometrial cancer cell line Ishikawa was used in the study. Cells were incubated with estradiol, followed by RNA isolation. We used the microarray method to identify estradiol-induced miRNAs in EC cells. Then, we analyzed tissues derived from 45 patients (18 EC1, 12 EC2, and 15 healthy endometrium (HE)) that were cut using the Laser Capture Microdissection method. The expression of selected miRNAs and their targets was assessed using the RT-qPCR method. Ishikawa cells were transfected with miRNA mimic, miRNA inhibitor (anti-miRNA), and their scrambled controls. We identified 66 estrogen-upregulated miRNAs in endometrial cancer cells. Out of them, miR-182 was upregulated in EC1 compared to HE. We found that miR-182 is an oncomiR in EC since its increased expression promoted the proliferation of EC cells and decreased expression of miR-182 was associated with the inhibition of cancer cell proliferation. Moreover, miR-182 inhibition upregulated SMAD4 expression. Our study allowed us to better understand the role of estrogen in the pathogenesis of EC.

## Introduction

Endometrial cancer (EC) is one of the most frequent cancers in women of perimenopausal age with growing incidence rates (Crosbie, Kitson et al. 2022). The first symptom is usually abnormal vaginal bleeding. There are no screening programs for early EC diagnosis (Crosbie, Kitson et al. 2022). Diagnosis is based on a transvaginal ultrasound scan and endometrial biopsy. Obesity is one of the most important risk factors of EC due to the estrogenic stimulation of endometrium unopposed by progesterone which leads to its proliferation (Crosbie, Kitson et al. 2022). Risk factors include other situations in which exposure to estrogens is prolonged, such as early menarche, late menopause, cycles without ovulations, and exogenous intake of estrogens (Nees, Heublein et al. 2022). Some genetic factors are also associated with the predisposition to endometrial cancer including Cowden syndrome or Lynch syndrome (Shai, Segev et al. 2014, Dominguez-Valentin, Sampson et al. 2020).

Endometrial cancer is divided into two types 1 and 2. EC1 is an estrogen-dependent EC that is characterized by a better outcome. Whereas type 2 is rare but associated with a worse prognosis (Henley, Ward et al. 2020). EC2 is responsible for 40% of deaths in the EC population even though it represents only 10-20% of EC cases (Setiawan, Yang et al. 2013). Recently proposed classification of EC is based on its molecular characteristics. There are four types of EC, ultramutated (POLE-mutant), hypermutated (mismatch repair deficient), copy number-high (p53-abnormal), and copy number-low, each characterized by different prognosis for patients (Crosbie, Kitson et al. 2022).

MiRNAs are small non-coding molecules that emerge as promising biomarkers of EC (Hutt, Tailor et al. 2019). They regulate the pathogenesis of different diseases by targeting the mRNAs of members of various cellular pathways (Grzywa, Klicka et al. 2019, Grzywa, Klicka et al. 2020). The expression of numerous miRNAs is dysregulated in EC tissues and their expression is correlated with patients’ outcomes (Klicka, Grzywa et al. 2022). Moreover, they regulate different cell signaling pathways in EC cells and thus they can inhibit (tumor suppressor miRNA) or promote cancer invasiveness and metastasis (oncomiRs) (Grzywa, Klicka et al. 2020, Klicka, Grzywa et al. 2021). Previous studies revealed that estrogens can induce expression of oncomiRs in breast cancer (Nagpal, Ahmad et al. 2013). Nonetheless, little is known regarding the role of the estrogens-miRNAs axis in EC. Hence, the aim of this study was to assess the role of the estrogen-induced miRNAs in EC.

## Materials and Methods

### Cell culture and treatment

Ishikawa cell line, estrogen and progesterone receptors-positive endometrial adenocarcinoma was used in this study. Cell line was purchased from Sigma-Aldrich. Cells were cultured in standard conditions (5% CO2, humified atmosphere, 37°C) using recommended Dulbecco’s Modified Eagle’s Medium (DMEM, Gibco, USA) containing 10% fetal bovine serum (FBS, Hyclone) according to the guidelines. To evaluate the role of estradiol in endometrial cancer in vitro, cells were incubated with 17β-estradiol 10^−8^ M (Sigma, USA) for 48 hours.

### RNA isolation, reverse transcription, and microarrays

RNA was isolated from cells incubated with estradiol and controls using RNeasy Mini Kit (Qiagen). Isolated RNA was eluted in 30 µl of ultrapure water and then stored in -80°C. The RNA purity and quantity were evaluated using NanoDrop 2000 spectrophotometer (Thermo Fisher Scientific, USA). cDNA was synthesized using Megaplex™ RT Primers, Human Pool Set v3.0 (Thermofisher) and TaqMan™ MicroRNA Reverse Transcription Kit (Thermofisher). Real-time PCR mix was prepared using TaqMan® Universal PCR Master Mix, No AmpErase UNG. The 100 µl aliquot was added to each port of TaqMan™ Array Human MicroRNA A+B Cards Set v3.0. The reactions were conducted using ViiA™ 7 Real-Time PCR System in universal cycling conditions (95°C/10 min, then 95°C/15 sec, 60°C/60 sec for 40 cycles). All reactions were performed according to the manufacturer’s protocols. Cycle threshold (Ct) values were used to analyze 754 human microRNAs. The expression of miRNAs selected in this step was subsequently assessed in EC tissues.

### Tissue specimens

In this study, we collected tissues of 30 patients diagnosed with endometrial cancer (EC) consisting of 18 type I EC and 12 type II EC, as well as 15 patients with healthy endometrium. Archival, formalin-fixed, paraffin-embedded (FFPE) tissues were obtained from the Department of Pathology, Medical Center of Postgraduate Education, Warsaw, Poland, and assessed there by a pathologist. Medical data such as a pathological description of collected tissues of endometrial cancer and healthy endometrium was received in anonymized form (Table 1).

**Table 1.** The histopathological and clinical data of patients.

| ID | Age | Type of EC/ HE | Grading | TNM |
| --- | --- | --- | --- | --- |
| 101 | 53 | Endometrioid EC (EC1) | G2 | pT3b N0 |
| 102 | 66 | Endometrioid EC (EC1) | G1 | pT1b N0 |
| 103 | 60 | Endometrioid EC (EC1) | G3 | pT1a N1 |
| 104 | 46 | Endometrioid EC (EC1) | G1 | pT1a N0 |
| 105 | 60 | Endometrioid EC (EC1) | G1 | pT2 N0 |
| 106 | 70 | Endometrioid EC (EC1) | G1 | pT1a N0 |
| 107 | 63 | Endometrioid EC (EC1) | G2 | pT2 N0 |
| 108 | 68 | Endometrioid EC (EC1) | G1 | pT1b Nx |
| 109 | 64 | Endometrioid EC (EC1) | G1 | pT3b N0 |
| 110 | 79 | Endometrioid EC (EC1) | G2 | pT1b N0 |
| 111 | 57 | Endometrioid EC (EC1) | G1 | pT1a N0 |
| 112 | 65 | Endometrioid EC (EC1) | G1 | pT3b N0 |
| 113 | 64 | Endometrioid EC (EC1) | G1 | pT1a N0 |
| 114 | 52 | Endometrioid EC (EC1) | G2 | pT2 N0 |
| 115 | 62 | Endometrioid EC (EC1) | G2 | pT3a N0 |
| 116 | 68 | Endometrioid EC (EC1) | G1 | pT1b N1 |
| 117 | 71 | Endometrioid EC (EC1) | G2 | pT1a N0 |
| 118 | 61 | Endometrioid EC (EC1) | G1 | pT1a N0 |
| 119 | 64 | Serous EC (EC2) | High grade | pT3a N0 |
| 120 | 84 | Serous EC (EC2) | n/a | pT1a N0 |
| 121 | 80 | Serous EC (EC2) | High grade | pT3b N1a |
| 122 | 79 | Serous EC (EC2) + Endometrioid EC (EC1) | High grade + G1 | pT3c N0 + pT1a N0 |
| 123 | 68 | Serous EC (EC2) | High grade | pT3b N0 |
| 124 | 64 | Serous EC (EC2) | High grade | pT1b N1 |
| 125 | 78 | Serous EC (EC2) | High grade | pT1b Nx |
| 126 | 73 | Serous EC (EC2) | n/a | pT3b N1 |
| 127 | 84 | Serous EC (EC2) | High grade | pT1a N0 |
| 128 | 69 | Serous EC (EC2) | High grade | pT2 N1a |
| 129 | 78 | Serous EC (EC2) | High grade | pT3b N2 |
| 130 | 66 | Serous EC (EC2) | High grade | pT3b N2 |
| 131 | 23 | HE | n/a | n/a |
| 132 | 46 | HE | n/a | n/a |
| 133 | 44 | HE | n/a | n/a |
| 134 | 41 | HE | n/a | n/a |
| 135 | 49 | HE | n/a | n/a |
| 136 | 50 | HE | n/a | n/a |
| 137 | 48 | HE | n/a | n/a |
| 138 | 55 | HE | n/a | n/a |
| 139 | 37 | HE | n/a | n/a |
| 140 | 51 | HE | n/a | n/a |
| 141 | 46 | HE | n/a | n/a |
| 142 | 52 | HE | n/a | n/a |
| 143 | 51 | HE | n/a | n/a |
| 144 | 47 | HE | n/a | n/a |
| 145 | 50 | HE | n/a | n/a |

### Preparation of archival samples for laser capture microdissection (LCM)

All FFPE samples were cut to 10 µm sections using a microtome and mounted on glass slides (SuperFrost Ultra Plus, Menzel Gläser) with a drop of DNAse/RNAse-free water. To increase adherence of slides, samples were incubated for an hour in a fume hood at 56 °C. Next, mounted tissue slices were hematoxylin and eosin stained according to the standard protocol in a set of stains, alcohol solutions, and xylene. Slides were subjected to LCM immediately after staining.

### Laser capture microdissection

LCM was performed using Zeiss PALM MicroBeam, Germany. Fragments of endometrial cancer tissue or healthy endometrium were cut out from the whole slide to avoid bias resulting from surrounding non-glandular tissues. The cut-out regions were marked before by a pathologist. Approximately 10mm2 of each section was cut and catapulted to the cap with 20 µl of Digestion Buffer.

### RNA isolation from tissue, reverse transcription, and real-time quantitative polymerase chain reaction (RT-qPCR)

RNA from the dissected fragment of tissue was isolated using RNAesy Mini Kit (Qiagen) following the manufacturer’s protocols. RNA was eluted in 30 µl of ultrapure water and stored at -80°C until the next steps. The quality of extracted RNA was assessed using NanoDrop 2000 spectrophotometer (Thermo Fisher Scientific, USA). Then, reverse transcription was conducted using Mir-X™ miRNA FirstStrand Synthesis Kit. Real-time qPCR was performed with Power SYBR™ Green PCR Master Mix (Thermo Fisher Scientific) and primers of miR-15a, -16, -27b, -28-3p, -29c, -31#, -92a, -106a, -106b, -129, -129-3p, -130a, -140, -140-3p, -141, -150, -152, -182, -192, -198, -200c, -215, - 335, -496, - 517c, -590-3p, and -656 (Table 2). U6 expression was used as normalization between samples. All reactions was performed in triplicates using ViiA 7 Real-Time PCR System (Thermo Fisher, USA). The 2^-ΔΔCt^ method was used to assess relative expression in analyzed tissues using the mean Ct values of selected microRNAs and U6.

**Table 2.**
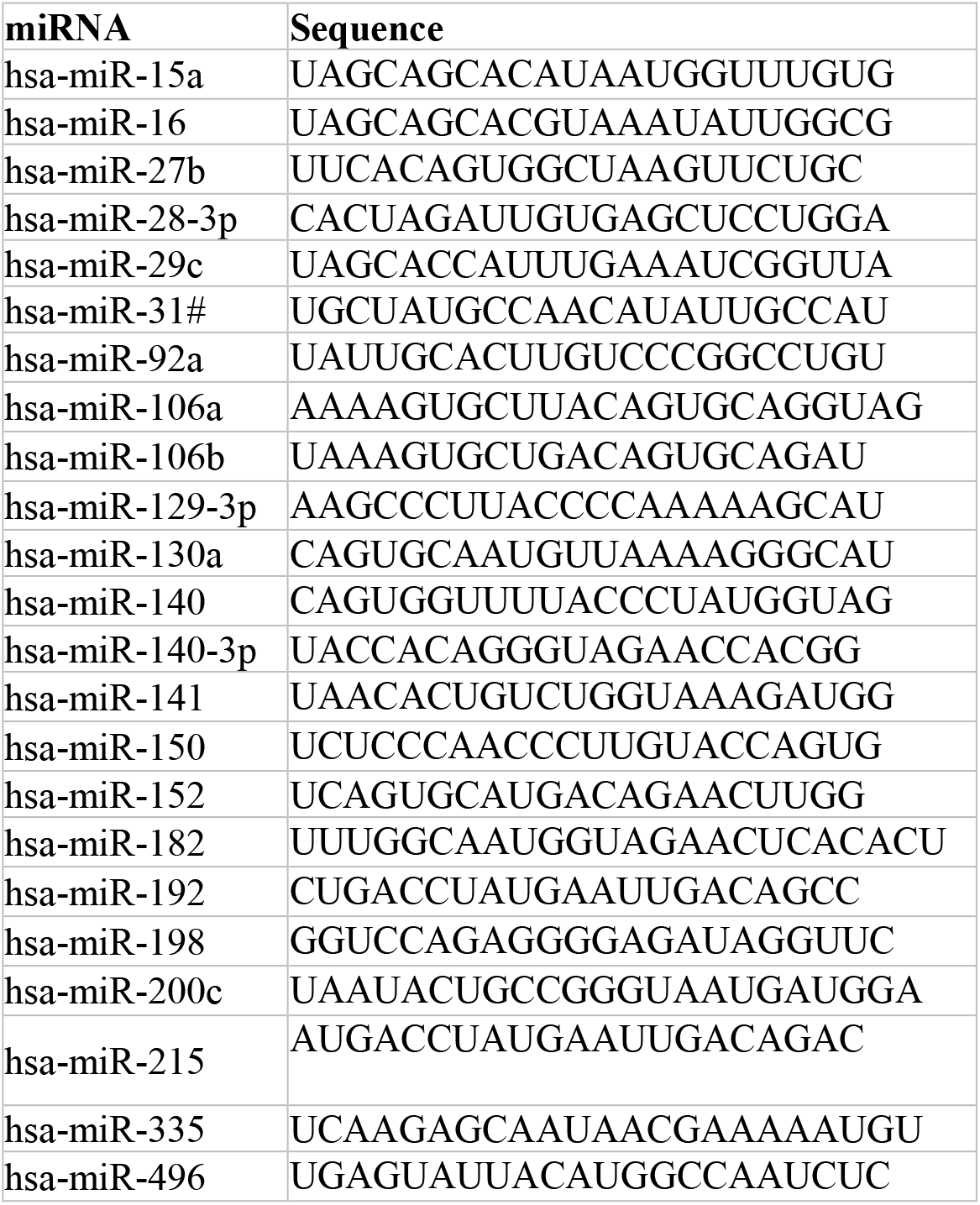
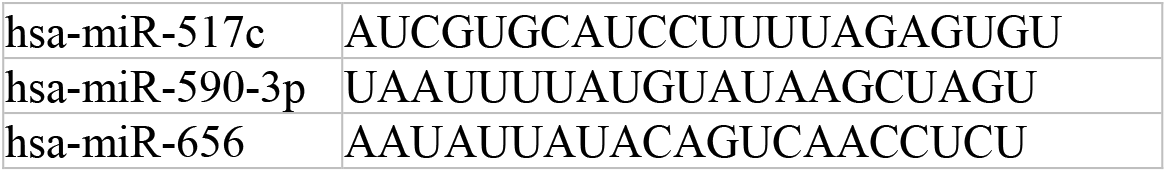
List of miRNAs’ primers sequences.

### Transfection

Transfection of Ishikawa was performed using DharmaFECT 1 siRNA Transfection Reagent according to the manufacturer’s protocol. Endometrial cancer cells were transfected with synthetic miR-mimic-182 and anti-miR-182, and their controls, miR-scrambled and anti-miR-scrambled obtained from Invitrogen™ mirVana™ (Thermo Fisher Scientific). The efficiency of transfection was assessed using the real-time qPCR method. Cells were then subjected to in vitro functional studies.

### Proliferation assay

For proliferation assay, 1 × 10^5^ cells/well were seeded in 12-well plates 24 h after transfection and were incubated for 48. Cells were then fixed with 4% paraformaldehyde, stained with 0.1% crystal violet, and photographed. The photos were analyzed using the ImageJ ColonyArea plugin (National Institutes of Health, Bethesda MD, USA) (Guzmán, Bagga et al. 2014).

### RNA isolation from cells, reverse transcription, and qRT-PCR

Total RNA was isolated from cells 24 h after transfection with miR-182 mimic, anti-miR-182 and their controls. Isolation was performed using RNAesy Mini Kit (Qiagen) according to the protocol. Then, RNA was reverse transcribed using Mir-X™ miRNA FirstStrand Synthesis Kit. Real-time qPCR was performed using Power SYBR™ Green PCR Master Mix (Thermo Fisher Scientific). The primer sequences of BCL6, CDH1-R, CDH2-R, COX2, Erα, ESRR1, EZH2, FN1, FOS, FOXA1, MARK-1, MCL-1, MMP9, NOTCH, PDL1, PFN, PTEN, RECK, SMAD4, SNAIL, TIM2, TIMP3, TP53, TWIST, VIM, ZBTB4 and ZEB1 are listed in Table 3. GAPDH was used as an endogenous control. The 2^-ΔΔCt^ method was used to calculate relative expression.

**Table 3.**
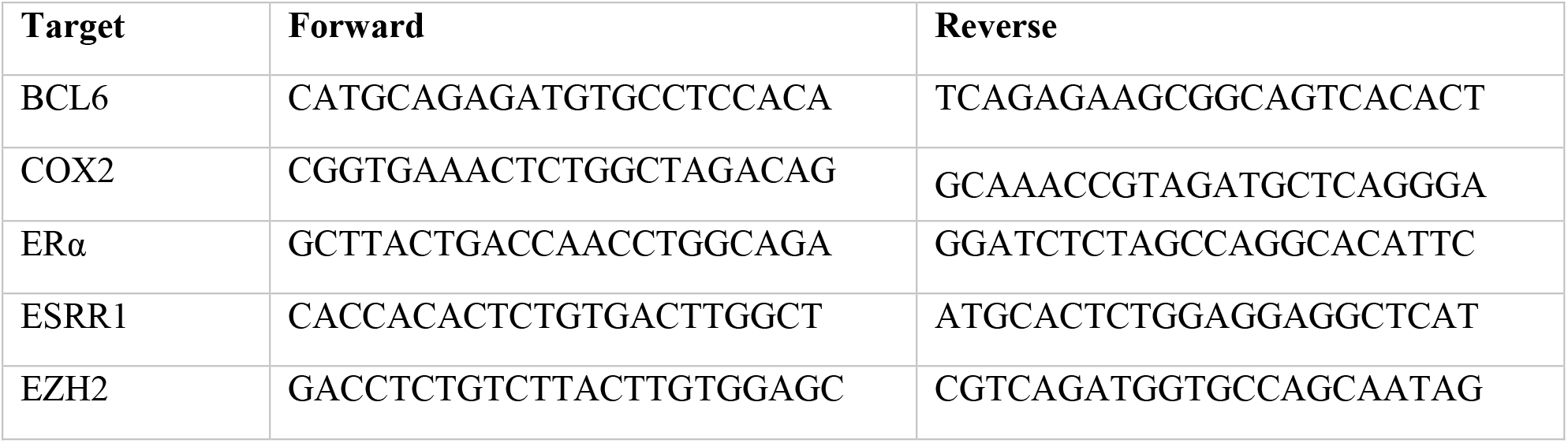
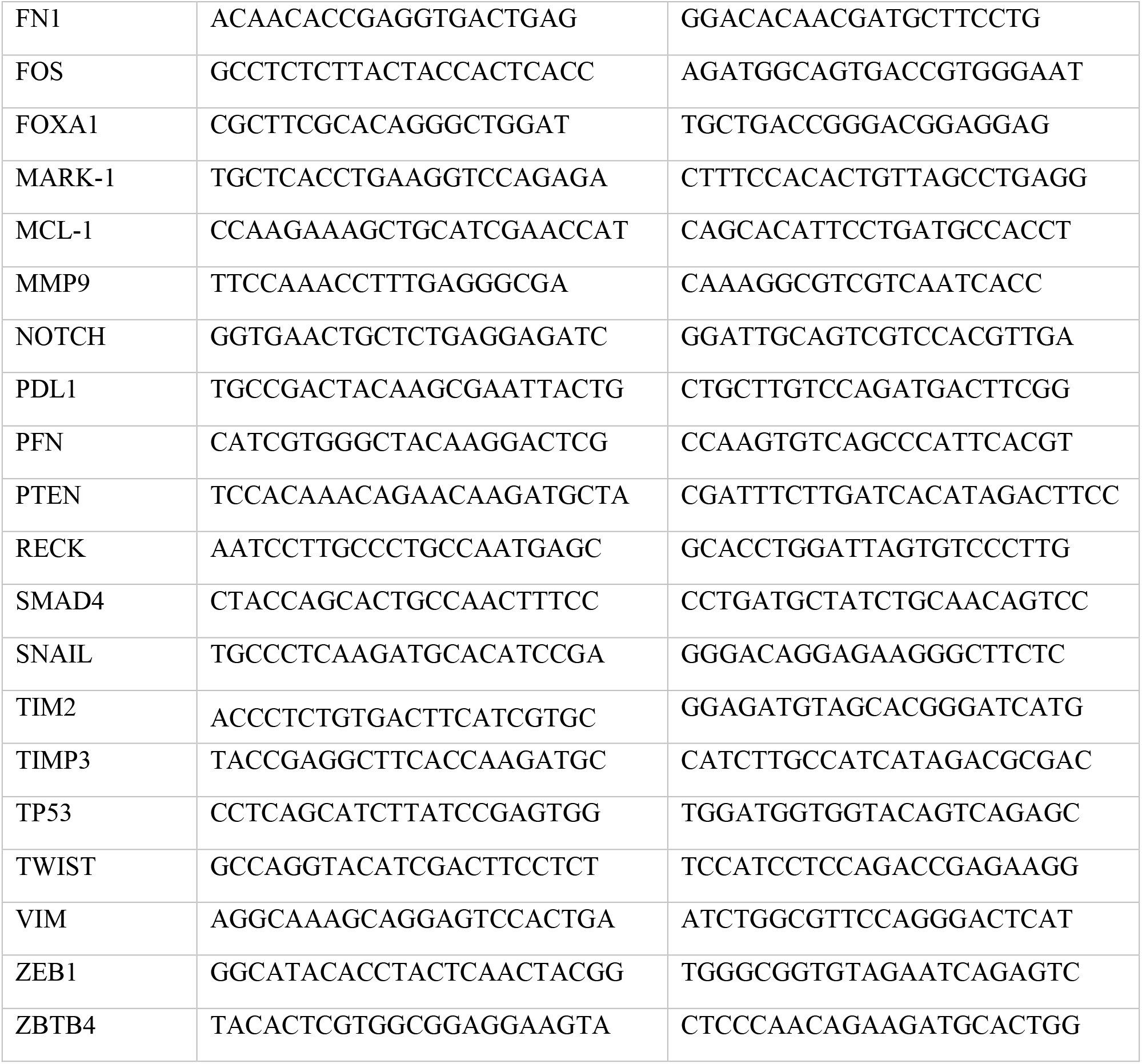
List of targets’ primers sequences.

### Data processing and analysis

All experiments were conducted according to the protocol in triplicates. GraphPad Prism 9.1 (GraphPad Software Inc.) was used for statistical analyses using the Kruskal-Wallis and paired t-test. A p-value lower than 0.05 was considered statistically significant. The figures were generated using Biorender. The Bioethical Committee of the Medical University of Warsaw declares (AKBE/78/2021) that the study is not considered a medical experiment under the Act of 5 December 1996 about the professions of doctor and dentist.

## Results

### Estradiol induces global changes in miRNA expression in endometrial cancer cells

To assess the role of the estrogen-miRNAs axis in EC (Fig. 1), we chose an Ishikawa endometrial cancer cell line that is characterized by the expression of estrogen receptors (Johnson, Maleki-Dizaji et al. 2007). These cells respond to estradiol stimulation with an increased proliferation (Supplementary Fig. 1). To determine a global profile of miRNA expression, we utilized the microarray method. We assessed the level of expression of a panel of 754 human miRNAs in Ishikawa cells that were incubated with estradiol for 48 hours. We detected 416 miRNAs in total and 66 miRNAs were upregulated in estradiol-treated Ishikawa cells. From them, we selected 26 miRNAs whose expression was most strongly upregulated by estradiol in EC cells (miR-15a, miR-16, miR-27b, miR-28-3p, miR-29c, miR-31#, miR-92a, miR-106a, miR-106b, miR-129-3p, miR-130a, miR-140, miR-140-3p, miR-141, miR-150, miR-152, miR-182, miR-192, miR-198, miR-200c, miR-215, miR-335, miR-496, miR-517c, miR-590-3p, and miR-656) (Table 4).

**Table 4.** Relative expression of selected miRNAs which expression was most strongly induced by estradiol in EC cells. Ishikawa EC cells were incubated with estradiol. The expression of miRNAs was assessed using microarray method. R = 2^-ΔΔCt^. The experiments were conducted in triplicates.

| miRNA | R1 | R2 | R3 |
| --- | --- | --- | --- |
| hsa-miR-15a | 2,467880839 | 1,057148936 | 1,521061477 |
| hsa-miR-16 | 1,038336523 | 1,13145572 | 1,579074587 |
| hsa-miR-27b | 1,608005906 | 1,102 | 1,883 |
| hsa-miR-28-3p | 1,256379062 | 1,216873893 | 2,101005552 |
| hsa-miR-29c | 1,260741828 | 1,121304206 | 2,170577526 |
| hsa-miR-31# | 2,321803776 | 6,03526599 | 5,412045038 |
| hsa-miR-92a | 1,029734961 | 1,019017726 | 1,615611437 |
| hsa-miR-106a | 1,086187165 | 1,195139224 | 1,31228322 |
| hsa-miR-106b | 1,675129604 | 1,929 | 1,225 |
| mmu-miR-129-3p | 2,95318864 | 12,136 | 58,895 |
| hsa-miR-130a | 1,524431298 | 1,020431937 | 2,056341445 |
| mmu-miR-140 | 1,835626692 | 1,06819722 | 1,622343006 |
| hsa-miR-140-3p | 1,252900716 | 1,072649364 | 2,269036844 |
| hsa-miR-141 | 1,0899575 | 1,191830424 | 2,280072983 |
| hsa-miR-150 | 4,820805576 | 1,522222892 | 4,039225975 |
| hsa-miR-152 | 2,786151833 | 1,033957689 | 1,513699148 |
| hsa-miR-182 | 1,441201201 | 1,412428573 | 3,600208353 |
| hsa-miR-192 | 4,017432428 | 1,989184912 | 2,086489757 |
| hsa-miR-198 | 4,311754235 | 2,155733029 | 4,783538936 |
| hsa-miR-200c | 1,5457141 | 1,455161307 | 2,19783108 |
| hsa-miR-215 | 1,483776178 | 2,503898738 | 1,083786101 |
| hsa-miR-335 | 2,846667552 | 2,760977007 | 2,631852852 |
| mmu-miR-496 | 3,134579781 | 7,444528801 | 4,503593272 |
| hsa-miR-517c | 3,610684897 | 2,193413999 | 3,810761257 |
| hsa-miR-590-3P | 2,809368102 | 16,273 | 15,902 |
| hsa-miR-656 | 3,3040759 | 2,133788754 | 3,65830464 |

**Fig. 1.**
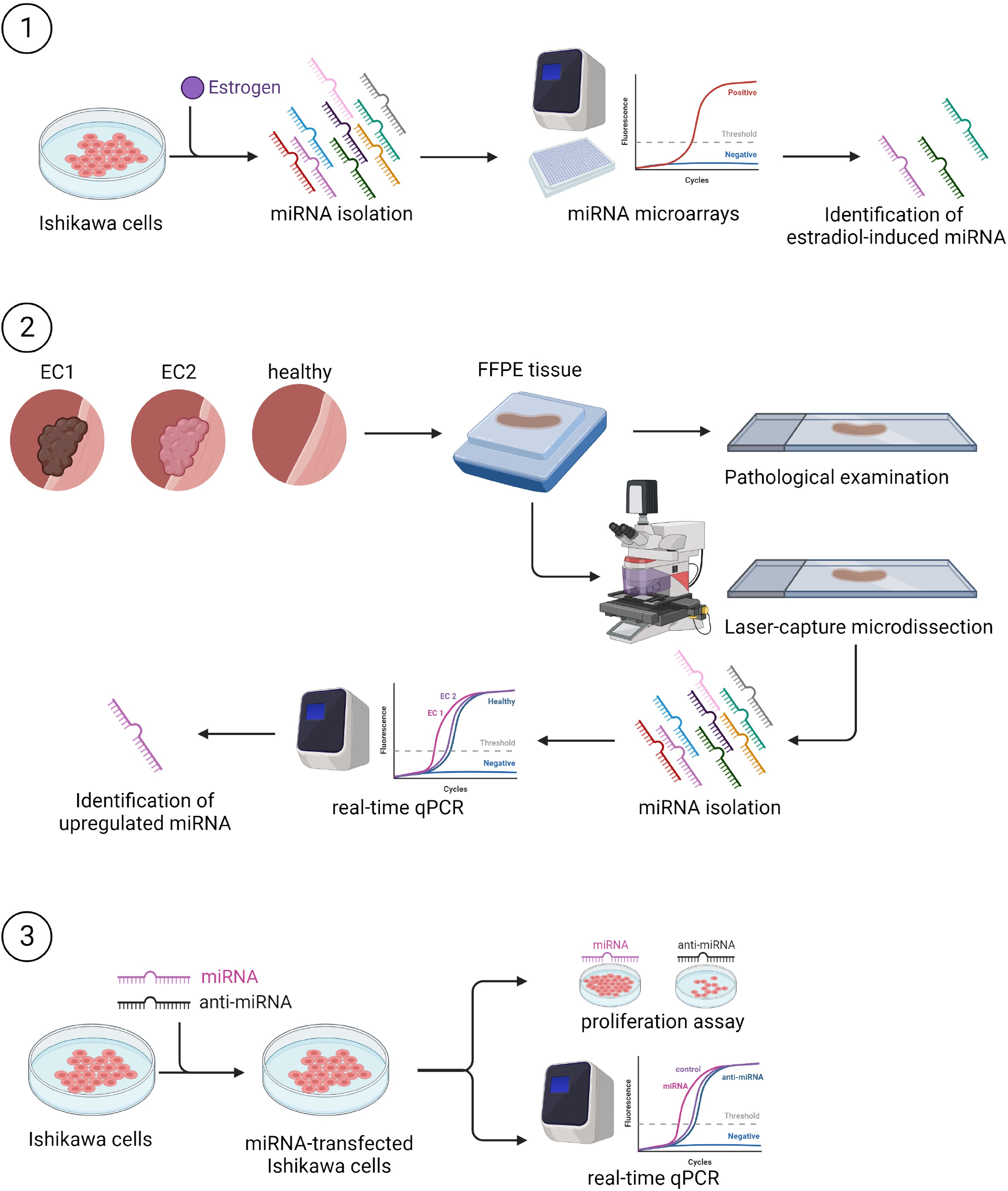
Study design diagram. (1) Ishikawa EC cells were incubated with estradiol. RNA was isolated. Estradiol-induced miRNAs were identified using the microarray method. (2) Selected regions of FFPE tissues of EC1, EC2 and HE were dissected. MiRNAs were isolated and their expression was assessed using RT-qPCR method. (3) Ishikawa EC cells were transfected with miRNA-mimic and its inhibitor and subjected to proliferation assay and RT-qPCR to find its target.

### MiR-182 is upregulated in endometrial cancer type I

To further confirm which miRNAs may be upregulated by estradiol in human EC, we analyzed tissues derived from 30 patients diagnosed with EC (18 EC1 and 12 EC2 samples) and 15 patients with healthy endometrium (HE) operated because of other reasons e.g., leiomyoma. We used the Laser Capture Microdissection method to isolate and selectively measure the expression of 26 miRNAs in cancer tissue. The results of the analyses revealed aberrant expression levels of three miRNAs. Notably, the expression of miR-182 was upregulated in EC1 compared to HE (Fig. 2a). MiR-27b expression was decreased in EC2 compared to HE (Fig. 2b) while the expression of miR-140 was decreased in both EC1 and EC2 compared to HE (Fig. 2c). There were no differences between the expression of miR-15a (Fig. 2d), miR-16 (Fig. 2e), miR-28-3p (Fig. 2f), miR-29c (Fig. 2g), miR-31# (Fig. 2h), miR-92a (Fig. 2i), miR-106a (Fig. 2j), miR-106b (Fig. 2k), miR-129 (Fig. 2l), miR-129-3p (Fig. 2m), miR-130a (Fig. 2n), miR-140-3p (Fig. 2o), miR-141 (Fig. 2p), miR-150 (Fig. 2q), miR-152 (Fig. 2r), miR-192 (Fig. 2s), miR-198 (Fig. 2t), miR-200c (Fig. 2u), miR-215 (Fig. 2v), miR-335 (Fig. 2w), miR-496 (Fig. 2x), miR-517c (Fig. 2y) and miR-656 (Fig. 2z) in EC1, EC2 and HE. The expression of miR-590-3p was too low to detect (data not shown). Thus, we selected miRNA-182 for further *in vitro* studies.

**Fig. 2.**
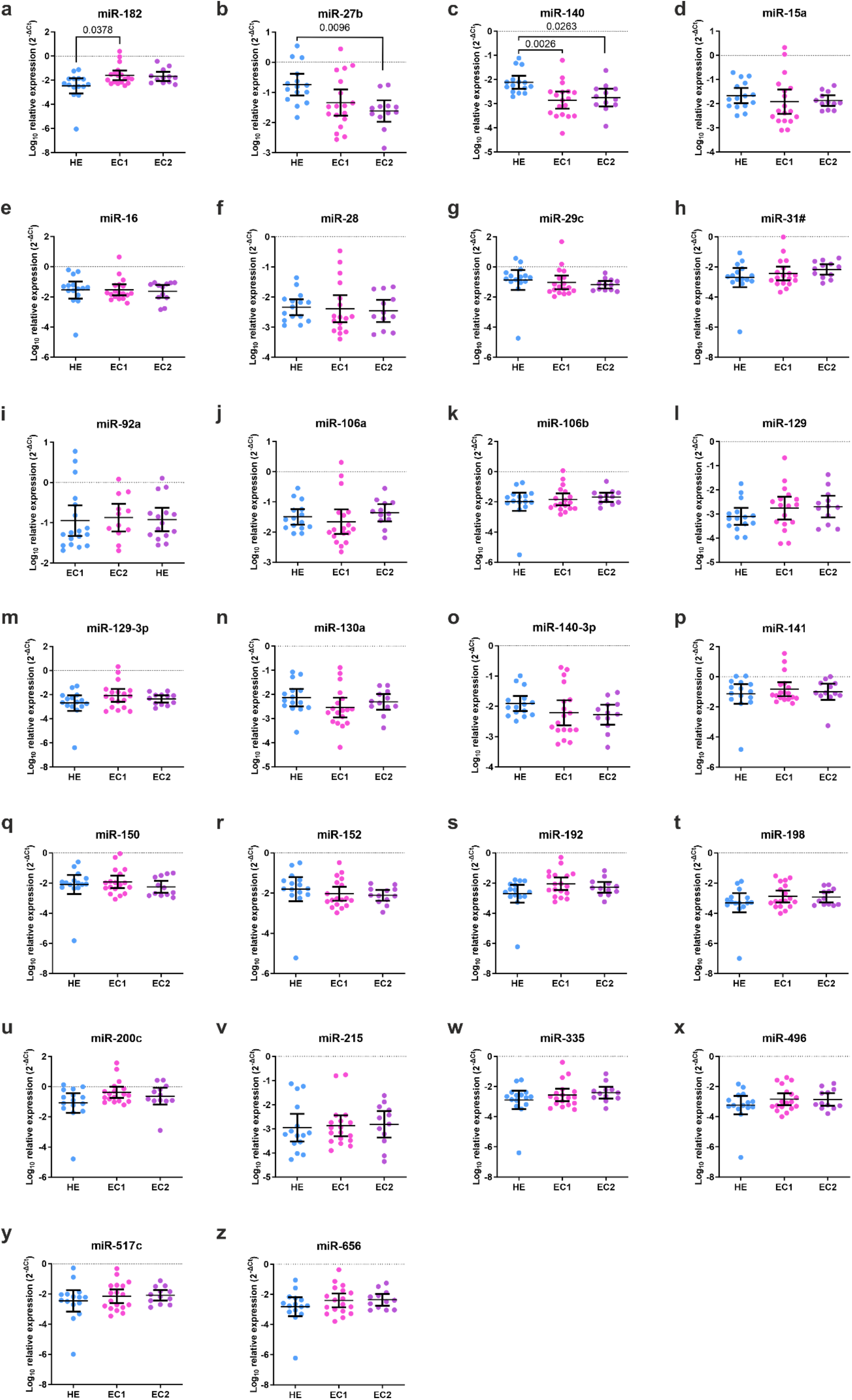
MiR-182 is upregulated in EC1 compared to HE. The expression of a) miR-182, b) miR-15a, c) miR-16, d) miR-27b, e) miR-28-3p, f) miR-29c, g) miR-31#, h) miR-92a, i) miR-106a, j) miR-106b, k) miR-129, l) miR-129-3p, m) miR-130a, n) miR-140, o) miR-140-3p, p) miR-141, q) miR-150, r) miR-152, s) miR-192, t) miR-198, u) miR-200c, v) miR-215, w) miR-335, x) miR-496, y) miR-517c, and z) miR-656 in dissected tissue of EC1, EC2 and HE. U6 was used as an endogenous control. *P*-values were calculated using the Kruskal-Wallis test. *p<0.05

### MiR-182 acts as an oncomiR in endometrial cancer

Our microarray results and increased expression of miR-182 in EC1 tissues compared to HE suggest miR-182 is induced by estrogens and may have a role in the pathogenesis of EC. To check that hypothesis we performed a proliferation assay. Ishikawa cells were transfected with a synthetic miR-182 mimic or miR-182 inhibitor (anti-miR-182) and their corresponding controls. We found that an increased level of miR-182 significantly enhanced the proliferation of Ishikawa cells (Fig. 3a), whereas inhibition of miR-182 resulted in an inhibition of the proliferation of EC cells (Fig. 3b). These studies revealed that miR-182 acts as an oncomiRs in EC.

**Fig. 3.**
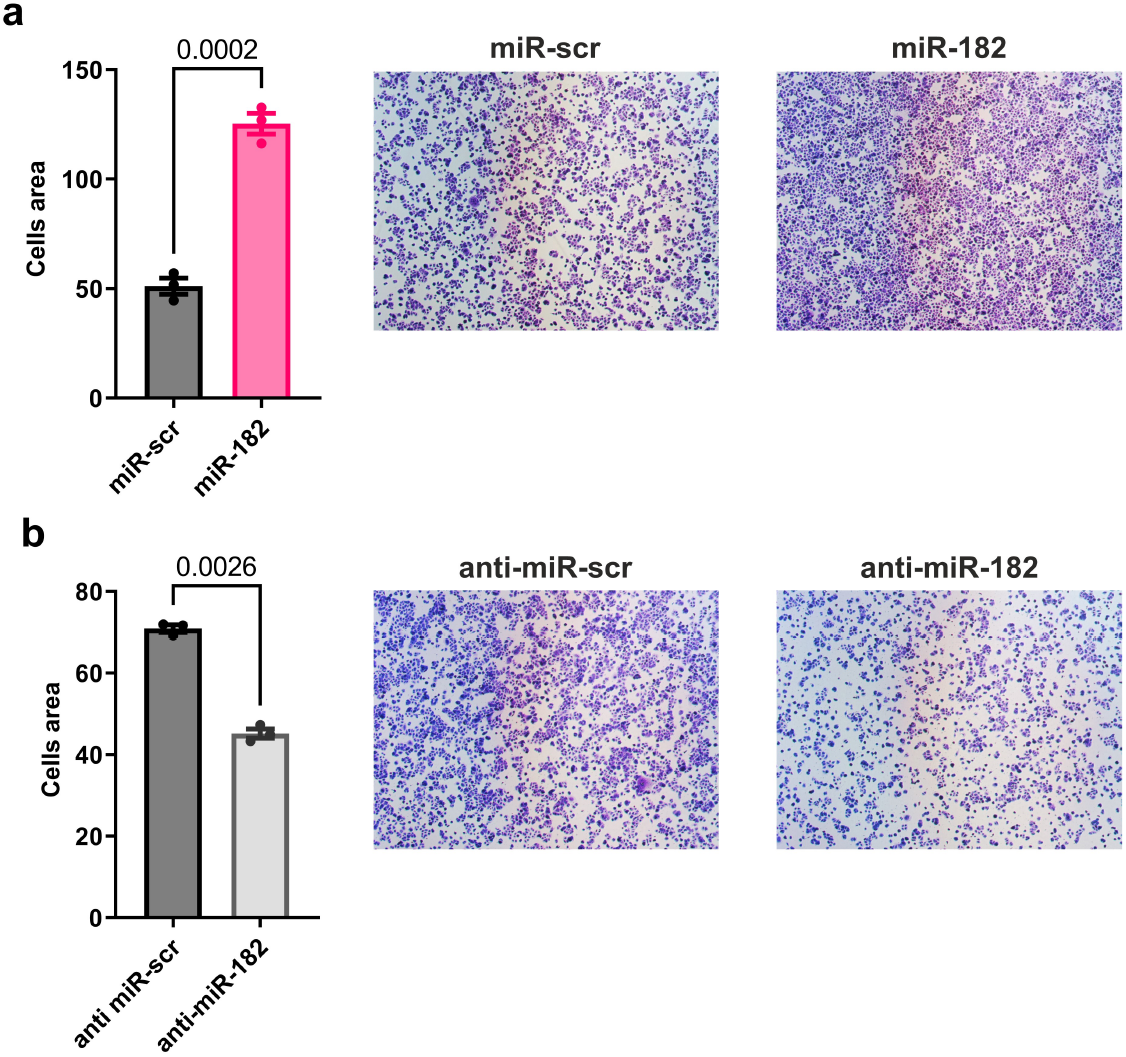
MiR-182 acts as oncomiRNA in EC cells. (a) The proliferation of mimic miR-182-transfected Ishikawa cells compared to control miR-scramble (miR-scr)-transfected cells and representative photos, p=0.0002. (b) The proliferation of anti-miR-182-transfected Ishikawa cells compared to control anti-miR-scr-transfected cells and representative photos, p=0.0026.

### Inhibition of miR-182 upregulates SMAD4 expression

Further, we sought to determine the mechanism of action of miRNA-182 in EC cells. We checked the role miRNA-182 plays in regulating the main oncogenic pathways in EC cells. For this purpose, Ishikawa cells were transfected with miR-182 mimic and its inhibitor. After 48 hours, the RNA was isolated, and then the expression of the targets selected based on the literature data was assessed. We determined the influence of miR-182 modulation on the expression of key proteins regulating epithelial-to-mesenchymal transition (EMT), apoptosis, and proliferation in EC (Grzywa, Klicka et al. 2020). We found that the inhibition of miR-182 in EC cells leads to the upregulated SMAD4 expression, which is an important member of TGFβ signaling (Fig. 4a) (Zhao, Mishra et al. 2018). We did not observe any influence of miR-182 on the expression of BCL2 (Fig. 4b), COX2 (Fig. 4c), ERα (Fig. 4d), ESRR1 (Fig. 4e), EZH2 (Fig. 4f), FN1 (Fig. 4g), FOS (Fig. 4h), FOXA1 (Fig. 4i), MARK1 (Fig. 4j), MCL1 (Fig. 4k), MMP9 (Fig. 4l), NOTCH (Fig. 4m), PDL1 (Fig. 4n), PFN (Fig. 4o), PTEN (Fig. 4p), RECK (Fig. 4q), TIM2 (Fig. 4r), TIMP3 (Fig. 4s), TP53 (Fig. 4t), TWIST (Fig. 4u), VIM (Fig. 4v), ZBTB4 (Fig. 4w), ZEB1 (Fig. 4y).

**Fig. 4.**
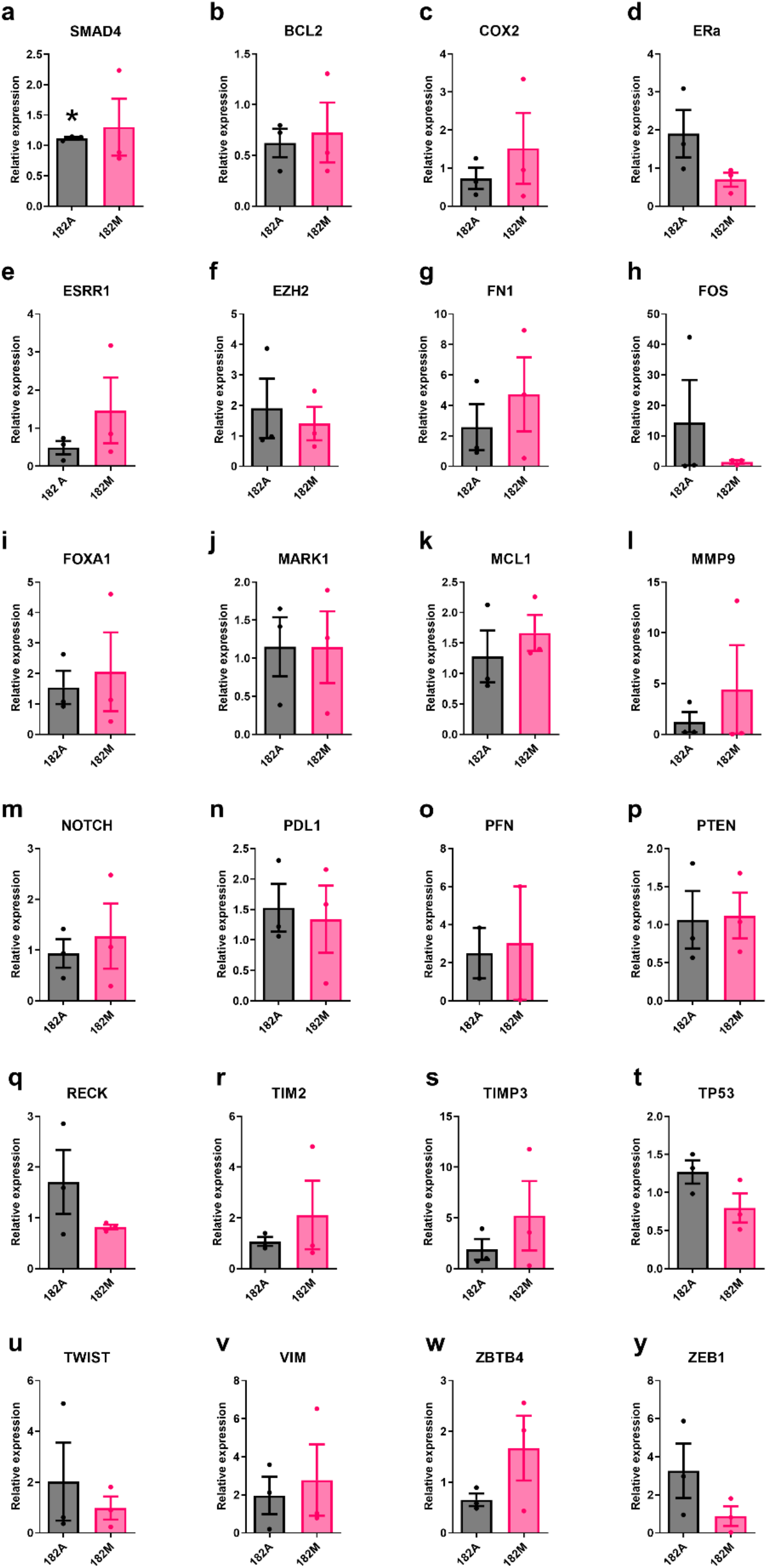
Inhibition of miR-182 upregulates SMAD4 expression. The relative expression of a) SMAD4, b) BCL2, c) COX2, d) ERα, e) ESRR1, f) EZH2, g) FN1, h) FOS, i) FOXA1, j) MARK1, k) MCL1, l) MMP9, m) NOTCH, n) PDL1, o) PFN, p) PTEN, q) RECK, r) TIM2, s) TIMP3, t) TP53, u) TWIST, v) VIM, w) ZBTB4, and y) ZEB1 in mimic miR-182 (182M) and anti-miR-182 (182a)-transfected Ishikawa cells compared to control miR-scramble-transfected Ishikawa cells. GAPDH was used as an endogenous control. *p=0.03.

## Discussion

This study demonstrated the global changes in miRNA expression induced by estradiol in EC cells. MiRNAs were described to have a crucial role in the pathogenesis of EC (Klicka, Grzywa et al. 2021). MiRNAs target mRNAs involved in the EMT process, cell cycle signaling, growth factor signaling, epigenetics, hormone signaling, and others (Klicka, Grzywa et al. 2021). Notably, the expression of various miRNAs was correlated with clinical parameters of patients with EC suggesting their potential as diagnostic and prognostic biomarkers (Klicka, Grzywa et al. 2021).

Some data in the literature already described the relationship between estrogens and miRNAs, but they were focused on breast cancer cells (Nagpal, Ahmad et al. 2013). Estrogen increases the proliferation of cancer cells, induces mostly oncomiRs, and represses tumor suppressor miRNAs (Cochrane, Cittelly et al. 2011). In the EC cell line rat model, estrogen was described to increase the expression of miR-181c and miR-200c (Zierau, Helle et al. 2018). These miRNAs target PTEN which regulates the PI3K-Akt axis, the key oncogenic pathway in EC cells (Chen, Zhang et al. 2018). Moreover, high expression of ERα was associated with overexpression of miR-200 (Indumati, Apurva et al. 2023). In this study, we detected 416 miRNAs, among which 66 were upregulated in Ishikawa EC cells incubated with estrogen. Out of them we chose 26 miRNAs and checked their expression in EC1, EC2 compared to HE (Table 4). The expression of miR-27b, miR-140, and miR-182 was changed in EC tissue.

Previous studies reported upregulated expression of miRNA-106a, miR-141, miR-182, miR-192, and miR-200c in EC human tissues. MiR-106a was described to act as an oncomiRNA promoting EC cell proliferation, invasion, and tumor growth by targeting BCL2L11 (Tang, Li et al. 2017). Upregulated expression of miR-141 in EC was associated with positive ER status, however, its expression is not statistically significantly correlated with clinical features (Dong, Si et al. 2015). Upregulated expression of miRNA-192 was associated with lymphovascular invasion (Lee, Ratner et al. 2014). MiR-200c expression was described to be upregulated in EC tissues however its role in EC is unclear (Klicka, Grzywa et al. 2021). The expression of miRNA-152 was downregulated in EC (Klicka, Grzywa et al. 2021). However, our study did not confirm these changes, excluding the role of these miRNAs as biomarkers of EC.

In our study, miR-182 was induced by estradiol in the EC cell line and upregulated in EC 1 compared to HE. MiR-182 was described to be upregulated in EC compared to HE tissue in both proliferative and secretory phases (Lee, Choi et al. 2012, Nishijima, Inoue et al. 2022) and in EC cell lines compared to human normal endometrial epithelial cells (Guo, Liao et al. 2013). Moreover, inhibition of miR-182 increased apoptosis of EC cells [18]. MiR-182 targets directly transcription elongation factor A-like 7 (TCEAL7). Inhibition of miR-182 decreased the proliferation of EC cells by downregulating NFκB-p65, cyclin D1, and c-Myc (Guo, Liao et al. 2013). Moreover, miR-182 directly binds to FOXO1 and FoxF2 and thus promotes proliferation, invasion, and migration (Fang, Sang et al. 2018, Yao, Kong et al. 2019). MiR-182 was described as a promising biomarker for the survival of EC patients (Wang, Zhong et al. 2015) as well as a possible therapeutic agent (Sameti, Tohidast et al. 2023). In our study, miR-182 increased the proliferation of EC cells which is consistent with literature data. Moreover, inhibition of miR-182 increased SMAD4 expression. The TGF-β/Smad pathway is one of the most important pathways in EC pathogenesis. These broad effects of miR-182 may be potentiated by its upregulation by estrogen in EC. SMAD4 is a known tumor suppressor in numerous cancer types (Zhao, Mishra et al. 2018). It interacts with key signaling pathways including MAPK, PI3K/Akt, and WNT/β-catenin (Zhao, Mishra et al. 2018). Thus, it influences the main processes in cancer cells, among them the EMT process in which cancer cells acquire the ability to migrate (Chen, Chiang et al. 2021). Downregulation of SMAD4 in EC causes repression of junction and adhesion complex genes leading to the increased motility of cancer cells (Lin, Kost et al. 2020). Moreover, lower expression of SMAD4 correlates with higher grading and worse prognosis of EC patients (Mhawech-Fauceglia, Kesterson et al. 2011). The TGF-β pathway members are described as targets of miRNAs in EC. SMAD4 is targeted by miR-27a-5p in EC which resulted in an inhibition of invasion and migration of EC cells (Che, Jian et al. 2020). Similarly, in other types of cancers, SMAD4 is a target of different miRNAs which promote cancer development (Zeng, Zhu et al. 2017, Zhou, Zhang et al. 2020). In ovarian cancer, SMAD4 is targeted by miR-145-5p and causes enhanced proliferation of cancer cells (Zhou, Zhang et al. 2020). It was reported that miR-205 targets SMAD4 in lung cancer resulting in increased growth of cancer cells *in vitro* and in *in vivo* studies (Zeng, Zhu et al. 2017).

The main limitation of the study is the small study group and lack of data concerning the outcome of patients, so the results of our study cannot be correlated with the patient’s prognosis. There is a need for further studies assessing the mechanism of action of estradiol-induced miRNAs in EC in vitro, as well as their association with clinical features of patients.

## Conclusions

In this study, we found that miR-182 is induced by estrogen in the EC cell line. Its expression was upregulated in tissue of EC 1 compared to healthy endometrium. *In vitro* studies revealed that miR-182 increased the proliferation of EC cells. Inhibition of miR-182 in EC resulted in increased expression of SMAD4. Additional studies are needed to assess the mechanism of action of miR-182, its role in EC in *in vivo* studies, and its potential clinical use as a biomarker.

## Acknowledgments

This study was funded by the National Centre of Science, grant number 2021/41/N/NZ5/02772 (K.K.). T.M.G. is supported by the National Centre of Science (2021/41/N/NZ6/02774) and the Foundation for Polish Science. The APC was funded by the Medical University of Warsaw. The funders had no role in study design, data collection and analysis, decision to publish, or preparation of the manuscript.

## Author Contributions

K.K. acquired funding, planned and conducted the experiments, analyzed the results, and wrote the manuscript. T.M.G. planned and conducted the experiments, analyzed the results, prepared figures, supervised the study, and wrote the manuscript. J.W. and J.O. performed collected and provided HE and EC samples and performed histopathological analysis of the samples. P.K.W. acquire funding and supervised the study. All authors have read and agreed to the published version of the manuscript.

## Data Availability Statement

The data that support the findings of this study are available from T.M.G. or P.K.W. upon reasonable request. Data from TCGA are publicly available.

## References

Che, X., F. Jian, C. Chen, C. Liu, G. Liu and W. Feng (2020). “PCOS serum-derived exosomal miR-27a-5p stimulates endometrial cancer cells migration and invasion.” J Mol Endocrinol 64(1): 1–12.

Chen, H. Y., Y. F. Chiang, J. S. Huang, T. C. Huang, Y. H. Shih, K. L. Wang, M. Ali, Y. H. Hong, T. M. Shieh and S. M. Hsia (2021). “Isoliquiritigenin Reverses Epithelial-Mesenchymal Transition Through Modulation of the TGF-β/Smad Signaling Pathway in Endometrial Cancer.” Cancers (Basel) 13(6).

Chen, R., M. Zhang, W. Liu, H. Chen, T. Cai, H. Xiong, X. Sheng, S. Liu, J. Peng, F. Wang, H. Chen, W. Lin, X. Xu, W. Zheng and Q. Jiang (2018). “Estrogen affects the negative feedback loop of PTENP1-miR200c to inhibit PTEN expression in the development of endometrioid endometrial carcinoma.” Cell Death Dis 10(1): 4.

Cochrane, D. R., D. M. Cittelly and J. K. Richer (2011). “Steroid receptors and microRNAs: relationships revealed.” Steroids 76(1-2): 1–10.

Crosbie, E. J., S. J. Kitson, J. N. McAlpine, A. Mukhopadhyay, M. E. Powell and N. Singh (2022). “Endometrial cancer.” Lancet 399(10333): 1412–1428.

Dominguez-Valentin, M., J. R. Sampson, T. T. Seppälä, S. W. Ten Broeke, J. P. Plazzer, S. Nakken, C. Engel, S. Aretz, M. A. Jenkins, L. Sunde, I. Bernstein, G. Capella, F. Balaguer, H. Thomas, D. G. Evans, J. Burn, M. Greenblatt, E. Hovig, W. H. de Vos Tot Nederveen Cappel, R. H. Sijmons, L. Bertario, M. G. Tibiletti, G. M. Cavestro, A. Lindblom, A. Della Valle, F. Lopez-Köstner, N. Gluck, L. H. Katz, K. Heinimann, C. A. Vaccaro, R. Büttner, H. Görgens, E. Holinski-Feder, M. Morak, S. Holzapfel, R. Hüneburg, M. V. Knebel Doeberitz, M. Loeffler, N. Rahner, H. K. Schackert, V. Steinke-Lange, W. Schmiegel, D. Vangala, K. Pylvänäinen, L. Renkonen-Sinisalo, J. L. Hopper, A. K. Win, R. W. Haile, N. M. Lindor, S. Gallinger, L. Le Marchand, P. A. Newcomb, J. C. Figueiredo, S. N. Thibodeau, K. Wadt, C. Therkildsen, H. Okkels, Z. Ketabi, L. Moreira, A. Sánchez, M. Serra-Burriel, M. Pineda, M. Navarro, I. Blanco, K. Green, F. Lalloo, E. J. Crosbie, J. Hill, O. G. Denton, I. M. Frayling, E. A. Rødland, H. Vasen, M. Mints, F. Neffa, P. Esperon, K. Alvarez, R. Kariv, G. Rosner, T. A. Pinero, M. L. Gonzalez, P. Kalfayan, D. Tjandra, I. M. Winship, F. Macrae, G. Möslein, J. P. Mecklin, M. Nielsen and P. Møller (2020). “Cancer risks by gene, age, and gender in 6350 carriers of pathogenic mismatch repair variants: findings from the Prospective Lynch Syndrome Database.” Genet Med 22(1): 15–25.

Dong, Y., J. W. Si, W. T. Li, L. Liang, J. Zhao, M. Zhou, D. Li and T. Li (2015). “miR-200a/miR-141 and miR-205 upregulation might be associated with hormone receptor status and prognosis in endometrial carcinomas.” Int J Clin Exp Pathol 8(3): 2864–2875.

Fang, Q., L. Sang and S. Du (2018). “Long noncoding RNA LINC00261 regulates endometrial carcinoma progression by modulating miRNA/FOXO1 expression.” Cell Biochem Funct 36(6): 323–330.

Grzywa, T. M., K. Klicka, B. Rak, D. Mehlich, F. Garbicz, G. Zieliński, M. Maksymowicz, E. Sajjad and P. K. Włodarski (2019). “Lineage-dependent role of miR-410-3p as oncomiR in gonadotroph and corticotroph pituitary adenomas or tumor suppressor miR in somatotroph adenomas via MAPK, PTEN/AKT, and STAT3 signaling pathways.” Endocrine 65(3): 646–655.

Grzywa, T. M., K. Klicka and P. K. Włodarski (2020). “Regulators at Every Step-How microRNAs Drive Tumor Cell Invasiveness and Metastasis.” Cancers (Basel) 12(12).

Guo, Y., Y. Liao, C. Jia, J. Ren, J. Wang and T. Li (2013). “MicroRNA-182 promotes tumor cell growth by targeting transcription elongation factor A-like 7 in endometrial carcinoma.” Cell Physiol Biochem 32(3): 581–590.

Guzmán, C., M. Bagga, A. Kaur, J. Westermarck and D. Abankwa (2014). “ColonyArea: an ImageJ plugin to automatically quantify colony formation in clonogenic assays.” PLoS One 9(3): e92444.

Henley, S. J., E. M. Ward, S. Scott, J. Ma, R. N. Anderson, A. U. Firth, C. C. Thomas, F. Islami, H. K. Weir, D. R. Lewis, R. L. Sherman, M. Wu, V. B. Benard, L. C. Richardson, A. Jemal, K. Cronin and B. A. Kohler (2020). “Annual report to the nation on the status of cancer, part I: National cancer statistics.” Cancer 126(10): 2225–2249.

Hutt, S., A. Tailor, P. Ellis, A. Michael, S. Butler-Manuel and J. Chatterjee (2019). “The role of biomarkers in endometrial cancer and hyperplasia: a literature review.” Acta Oncol 58(3): 342–352.

Indumati, S., B. Apurva, G. Gaurav, S. Nehakumari and V. Nishant (2023). “The Role of MicroRNAs in Development of Endometrial Cancer: A Literature Review.” J Reprod Infertil 24(3): 147–165.

Johnson, S. M., M. Maleki-Dizaji, J. A. Styles and I. N. H. White (2007). “Ishikawa cells exhibit differential gene expression profiles in response to oestradiol or 4-hydroxytamoxifen.” Endocrine-Related Cancer Endocr Relat Cancer 14(2): 337–350.

Klicka, K., T. M. Grzywa, A. Klinke, A. Mielniczuk, J. Wejman, J. Ostrowska, A. Gondek and P. K. Włodarski (2022). “Decreased expression of miR-23b is associated with poor survival of endometrial cancer patients.” Sci Rep 12(1): 18824.

Klicka, K., T. M. Grzywa, A. Klinke, A. Mielniczuk and P. K. Włodarski (2021). “The Role of miRNAs in the Regulation of Endometrial Cancer Invasiveness and Metastasis-A Systematic Review.” Cancers (Basel) 13(14).

Klicka, K., T. M. Grzywa, A. Klinke, A. Mielniczuk and P. K. Włodarski (2021). “The Role of miRNAs in the Regulation of Endometrial Cancer Invasiveness and Metastasis—A Systematic Review.” Cancers 13(14): 3393.

Lee, H., H. J. Choi, C. S. Kang, H. J. Lee, W. S. Lee and C. S. Park (2012). “Expression of miRNAs and PTEN in endometrial specimens ranging from histologically normal to hyperplasia and endometrial adenocarcinoma.” Mod Pathol 25(11): 1508–1515.

Lee, L. J., E. Ratner, M. Uduman, K. Winter, M. Boeke, K. M. Greven, S. King, T. W. Burke, K. Underhill, H. Kim, R. J. Boulware, H. Yu, V. Parkash, L. Lu, D. Gaffney, A. P. Dicker and J. Weidhaas (2014). “The KRAS-Variant and miRNA Expression in RTOG Endometrial Cancer Clinical Trials 9708 and 9905.” PLOS ONE 9(4): e94167.

Lin, L.-L., E. R. Kost, C.-L. Lin, P. Valente, C.-M. Wang, M. G. Kolonin, A. C. Daquinag, X. Tan, N. Lucio, C.-N. Hung, C.-P. Wang, N. B. Kirma and T. H. M. Huang (2020). “PAI-1-Dependent Inactivation of SMAD4-Modulated Junction and Adhesion Complex in Obese Endometrial Cancer.” Cell Reports 33(2).

Mhawech-Fauceglia, P., J. Kesterson, D. Wang, S. Akers, N. C. DuPont, K. Clark, S. Lele and S. Liu (2011). “Expression and clinical significance of the transforming growth factor-β signalling pathway in endometrial cancer.” Histopathology 59(1): 63–72.

Nagpal, N., H. M. Ahmad, B. Molparia and R. Kulshreshtha (2013). “MicroRNA-191, an estrogen-responsive microRNA, functions as an oncogenic regulator in human breast cancer.” Carcinogenesis 34(8): 1889–1899.

Nees, L. K., S. Heublein, S. Steinmacher, I. Juhasz-Böss, S. Brucker, C. B. Tempfer and M. Wallwiener (2022). “Endometrial hyperplasia as a risk factor of endometrial cancer.” Arch Gynecol Obstet 306(2): 407–421.

Nishijima, Y., N. Inoue, A. Iwase, H. Yokoo and M. Saio (2022). “MicroRNA 182, 183, 200a, and 200b exhibit strong correlations but no involvement in PTEN protein regulation in uterine endometrial carcinoma.” Pathol Res Pract 236: 153986.

Sameti, P., M. Tohidast, M. Amini, S. Z. Bahojb Mahdavi, S. Najafi and A. Mokhtarzadeh (2023). “The emerging role of MicroRNA-182 in tumorigenesis; a promising therapeutic target.” Cancer Cell Int 23(1): 134.

Setiawan, V. W., H. P. Yang, M. C. Pike, S. E. McCann, H. Yu, Y. B. Xiang, A. Wolk, N. Wentzensen, N. S. Weiss, P. M. Webb, P. A. van den Brandt, K. van de Vijver, P. J. Thompson, B. L. Strom, A. B. Spurdle, R. A. Soslow, X. O. Shu, C. Schairer, C. Sacerdote, T. E. Rohan, K. Robien, H. A. Risch, F. Ricceri, T. R. Rebbeck, R. Rastogi, J. Prescott, S. Polidoro, Y. Park, S. H. Olson, K. B. Moysich, A. B. Miller, M. L. McCullough, R. K. Matsuno, A. M. Magliocco, G. Lurie, L. Lu, J. Lissowska, X. Liang, J. V. Lacey, Jr., L. N. Kolonel, B. E. Henderson, S. E. Hankinson, N. Håkansson, M. T. Goodman, M. M. Gaudet, M. Garcia-Closas, C. M. Friedenreich, J. L. Freudenheim, J. Doherty, I. De Vivo, K. S. Courneya, L. S. Cook, C. Chen, J. R. Cerhan, H. Cai, L. A. Brinton, L. Bernstein, K. E. Anderson, H. Anton-Culver, L. J. Schouten and P. L. Horn-Ross (2013). “Type I and II endometrial cancers: have they different risk factors?” J Clin Oncol 31(20): 2607–2618.

Shai, A., Y. Segev and S. A. Narod (2014). “Genetics of endometrial cancer.” Fam Cancer 13(3): 499–505.

Tang, W., J. Li, H. Liu, F. Zhou and M. Liu (2017). “MiR-106a promotes tumor growth, migration, and invasion by targeting BCL2L11 in human endometrial adenocarcinoma.” Am J Transl Res 9(11): 4984–4993.

Wang, F., S. Zhong, H. Zhang, W. Zhang, H. Zhang, X. Wu and B. Chen (2015). “Prognostic Value of MicroRNA-182 in Cancers: A Meta-Analysis.” Dis Markers 2015: 482146.

Yao, H., F. Kong and Y. Zhou (2019). “MiR-182 promotes cell proliferation, migration and invasion by targeting FoxF2 in endometrial carcinoma cells.” Int J Clin Exp Pathol 12(4): 1248–1259.

Zeng, Y., J. Zhu, D. Shen, H. Qin, Z. Lei, W. Li, Z. Liu and J. A. Huang (2017). “MicroRNA-205 targets SMAD4 in non-small cell lung cancer and promotes lung cancer cell growth in vitro and in vivo.” Oncotarget 8(19): 30817–30829.

Zhao, M., L. Mishra and C. X. Deng (2018). “The role of TGF-β/SMAD4 signaling in cancer.” Int J Biol Sci 14(2): 111–123.

Zhou, J., X. Zhang, W. Li and Y. Chen (2020). “MicroRNA-145-5p regulates the proliferation of epithelial ovarian cancer cells via targeting SMAD4.” J Ovarian Res 13(1): 54.

Zierau, O., J. Helle, S. Schadyew, Y. Morgenroth, M. Bentler, A. Hennig, S. Chittur, M. Tenniswood and G. Kretzschmar (2018). “Role of miR-203 in estrogen receptor-mediated signaling in the rat uterus and endometrial carcinoma.” J Cell Biochem 119(7): 5359–5372.

